# Unsupervised learning of structural variability in cryo-EM data using normal mode analysis of deformable atomic models

**DOI:** 10.1101/2025.07.27.666683

**Authors:** Youssef Nashed, Julien Martel, Ariana Peck, Axel Levy, Huanghao Mai, Gordon Wetzstein, Nina Miolane, Daniel Ratner, Frédéric Poitevin

## Abstract

Cryogenic electron microscopy (cryo-EM) has emerged as the method of choice to characterize the structural variability of biomolecules at near-atomic resolution. We present a reconstruction approach that eliminates the need for post-hoc atomic model fitting in 3D maps by deforming a given atomic model along its normal modes directly against the 2D data. This end-to-end approach inherently reduces the risk of error propagation while increasing interpretability of resulting structural ensembles.

---

Understanding the structural variability of molecules in atomic detail is critical for drug and biomolecular design [1, 2]. Cryogenic electron microscopy (cryo-EM) has emerged as a promising method toward that goal [3] since it produces large numbers of 2D images of the same molecule in unknown conformations, containing information that enables 3D reconstructions at ever improving resolutions [4], which in turn enables fitting atomic models with increasing reliability [5]. While the laborious nature of that siloed two-step process [6] can be automated [7], structural variability is typically studied at the 3D map level which is not readily interpretable in terms of molecular mechanisms [8]. It would be desirable to build atomic models directly from 2D images to resolve this interpretability issue [9].

However, the high-dimensionality and rugged nature of molecular energy landscapes make them notoriously difficult to explore [10], a requirement to build realistic atomic models. To alleviate this difficulty, existing methods reduce the problem size by coarse-graining molecules and either treating them as point clouds [11, 12] or elastic network models (ENM) [13]. Normal mode analysis (NMA) [14] of ENMs permits reducing the dimensionality of the energy landscape to only its lowest frequency normal modes, which often encode functionally relevant motions [15–22]. Jin et al. first incorporated this approach into cryo-EM reconstruction, where coordinates along pre-computed normal modes are estimated for each image using a variant of the expectation-maximization (EM) algorithm [23]. While the EM algorithm guarantees reconstruction accuracy, it comes at the expense of scalability [24]. To remedy this, Hamitouche and Jonic train a neural network on an EM-labeled subset of the dataset and extrapolate to the remainder [25].

In order to build atomic models directly from 2D images without coarse-graining or suffering from the suboptimal trade-off typical of supervised approaches, an efficient implementation of all-atom model deformation coupled with a scalable and robust approach to learn the deformation is warranted. Here we introduce `aNiMAte`, a selfsupervised deep learning solution implemented as an auto-encoder (see Methods): images are encoded as normal mode coordinates subsequently decoded into synthetic images with a differentiable cryo-EM simulator (Fig. 1A). The implementation in the decoder of both molecular deformation from an initial atomic model and image formation from a deformed model has been optimized to efficiently tackle molecules of arbitrary sizes. We compute a per-atom self-consistency (Δ) score by training two models on two independent subsets of images. Atoms with a low self-consistency (high Δ) score are discarded from the atomic model. To visualize the structural variability in the cryo-EM dataset, principal component analysis (PCA) of the learned normal mode coordinates is carried out, highlighting the major types of motions observed in the dataset directly in the space of atomic models (Fig. 1C). We discover biologically relevant modes of motion in two publicly available experimental datasets of the pre-catalytic spliceosome [26] and the ribosome [27]. Importantly, sampling conformations along non-linear trajectories in the normal mode space leads to interesting and non-linear trajectories in atom coordinate space.

**Fig. 1.**
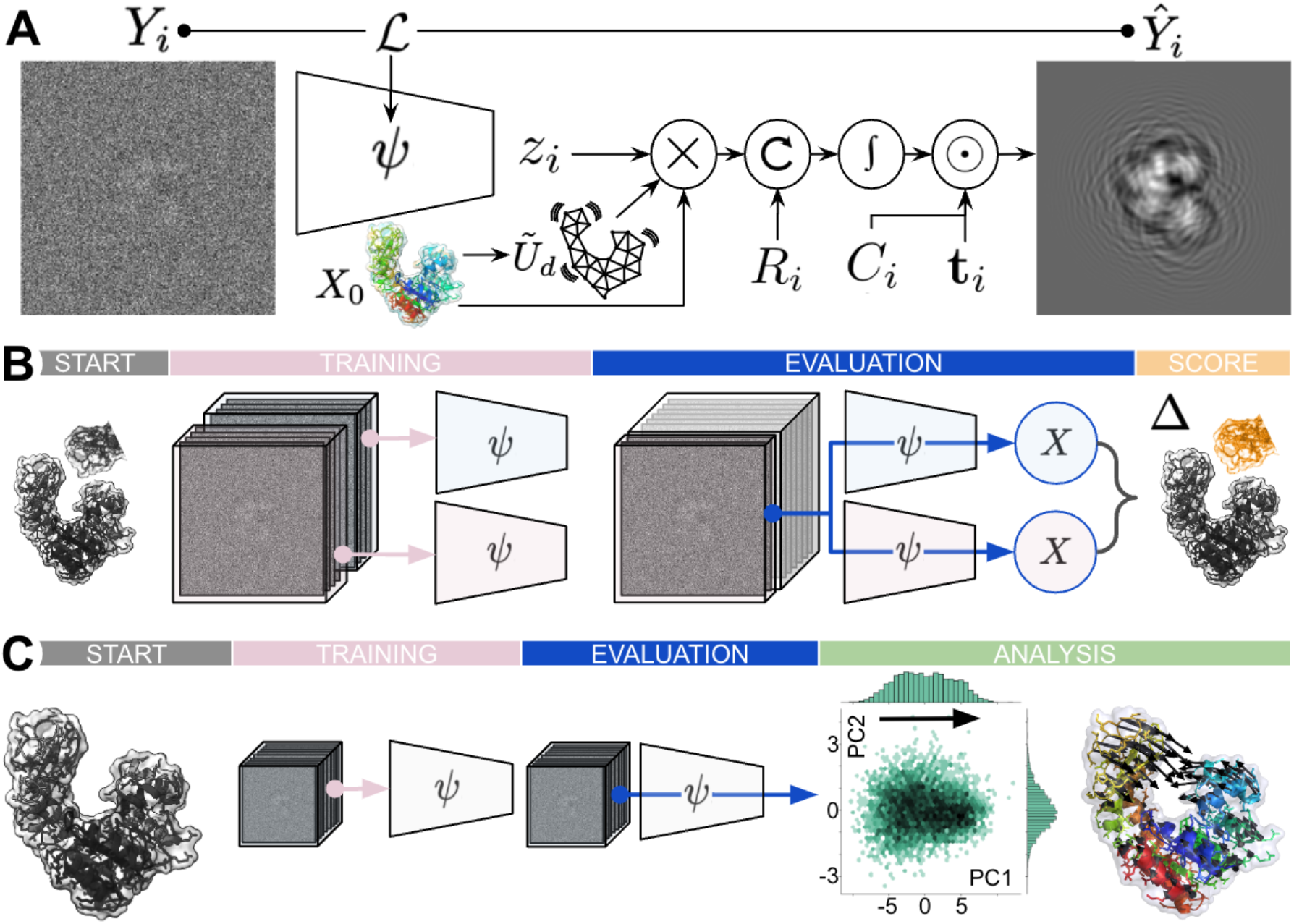
(A) Overview of aNiMAte. The encoder, parameterized by *ψ*, maps an image *Y*_*i*_ to the normal mode coordinates *z*_*i*_. The decoder first deforms an initial atomic model *X*_0_ along its normal modes 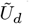 using the variables *z*_*i*_ learned by the encoder. The resulting deformed atomic model is rotated using a known orientation *R*_*i*_, and evaluated using the superposition of the electron form factors of the atoms at the desired pixel positions. Given per-image translation **t**_*i*_ and contrast transfer function (CTF) parameters *C*_*i*_, a di!erentiable physics-based simulator generates *Ŷ*_*i*_, a noise-free estimation of *Y*_*i*_. The input and output images are compared with the cross correlation loss ℒ, which is minimized for *ω*. **(B) Atomic model scoring**. Two encoders are trained independently on two halves of the dataset, using the same initial atomic model. Each encoder is then evaluated on a subset of the data and a Δ score is assigned to each atom, enabling discarding atoms with threshold-exceeding scores (in orange on the figure). **(C) Structural Variability Analysis**. The Δ-reduced atomic model is used to retrain a new encoder on the full dataset, or again to train two encoders on the two halves to produce uncertainties for the predictions. Principal Component Analysis over resulting *z*_*i*_ yields a conformation latent space readily interpretable with atomistic trajectories.

The procedure described in Fig. 1B was first applied on the atomic models deposited with their corresponding datasets, and atoms with a Δscore higher than 10Å were discarded (see Fig.S1). For both molecules, the core atoms display a Δscore below 1Å. The head, U2 region and foot of the spliceosome, and the head, L1 stalk and central protuberance of the ribosomes are typically associated with higher Δ scores, up to 3Å. Training on the whole dataset as illustrated in Fig. 1C yields a distribution over 16 normal mode coordinates that can be further reduced through principal component analysis for visualization purposes. For each molecule, at most two components are needed to explain 75% of the variance in both datasets (Fig. 2B). Selected points along trajectories in the space of the first two principal components are used to generate corresponding atomic models (Fig. 2C).

**Fig. 2.**
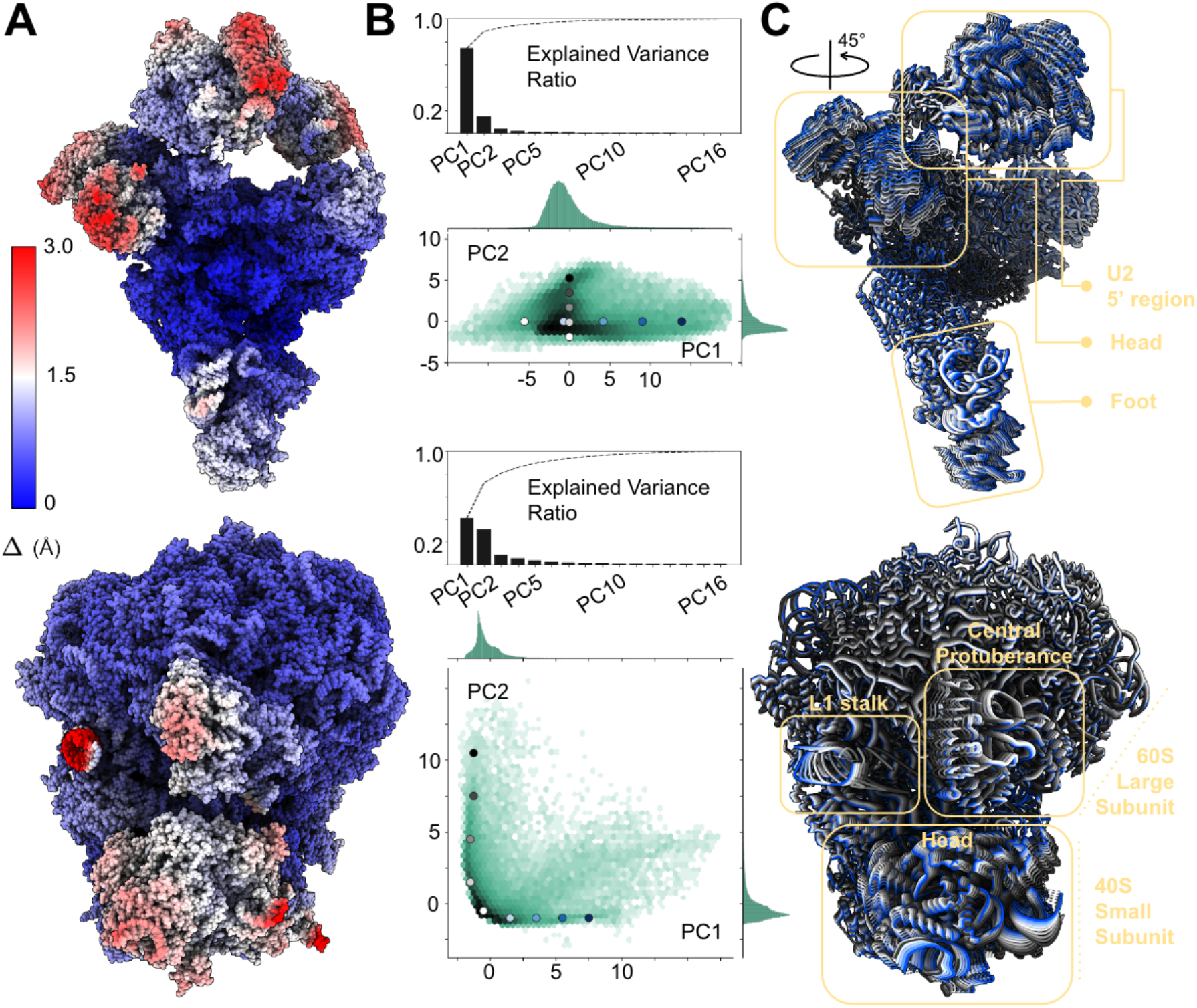
Results on experiment datasets: top - EMPIAR 10180 (spliceosome); bottom - EMPIAR 10028 (ribosome). The atomic model used to train `aNiMAte` is shown colored with its atomic model score Δ **(A)**. Principal Component Analysis of the learned normal mode coordinates results in a 2D space (PC1, PC2) explaining more than 75% of the dataset variability **(B)** illustrated with representative atomic models traversing it **(C)**.

On the same splicesome dataset, previous analysis explained 30% of structural variability at the 3D maps level using the first two principal components, and suggested that the head and 5’ region of the U2 particle moved independently [28, 29]. Here, we find that 74% of the variability in the data can be explained by a concerted motion of these two regions (Fig. 2B+C-top). Strikingly, this motion is mostly explained by the lowest frequency normal mode computed from the ENM of the Δ -reduced atomic model (Fig.S2A), even though independent domain motion can be represented using a combination of the 16 first normal modes.

For the ribosome dataset, `aNiMAte` scores the region surrounding the peptydil transferase center with Δ score beyond 1.5Å (Fig. 2A-bottom), in agreement with previously described local resolution maps [27]. Most of the variability in the dataset (Fig. 2B-bottom) is supported by two types of motions corresponding to ribosomal head swivel or to a concerted motion of the central protuberance and L1 stalk of the large subunits (Fig. 2A-bottom right) which are hallmarks of the ribosome translocation mechanism [30] and agree qualitatively with structural variability analysis done at the 3D maps level [28, 29] while providing an unprecedented level of mechanistic detail.

Comparing the variances of the predicted conformations along each normal mode with their ENM eigenvalues, we observe that these quantities are correlated (Fig.2S). This supports the assumption that, under the constraint of using *N* normal modes, the least amount of information is lost when reducing motion to the *N* modes of lowest energy. Interestingly, the motions we analyzed recruit non-trivial normal modes in addition to the lowest-frequency mode, highlighting how the deformation prior brought by NMA can be reweighted by the information in the cryo-EM data.

`aNiMAte` opens an avenue for the analysis of structural variability in cryo-EM datasets from 2D images without resorting to coarse-graining or supervision, thus enabling direct interpretation of molecular dynamics at the atomic level. We are hopeful that future work building on the method presented here will address possible limitations, for example by extending to non-linear deformation models [31], joint pose estimation, or compositional variability.

## Methods

### Image Formation Model

In single particle cryo-EM, probing electrons interact with the electrostatic potential created by molecules embedded in a thin layer of vitreous ice.

#### Elastic Network Models, Normal Mode Analysis and Coarse-Graining

The conformational space of the molecular structure is 3*M* dimensional, noting *M* its number of constitutive atoms ∼ 10^2^ − 10^5^. Efficient sampling of this very high-dimensional space to uncover the lower-dimensional subspace defined by a handful of collective variables (the *conformational space* 𝒱) has been an area of intense research for many decades [32–34]. A popular and efficient option involves performing a harmonic approximation of the energy landscape around a chosen conformation [14, 35] by representing the molecule with an elastic network model (ENM) [13, 36]. Noting *X* = {**r**_*j*_}_*j*=1,*M*_ the atomic Cartesian coordinatesof the molecule, the potential energy *E* of the ENM is further approximated with a second-order Taylor approximation around an initial atomic model *X*_0_ and using *H* the Hessian matrix of *E* in *X*_0_:

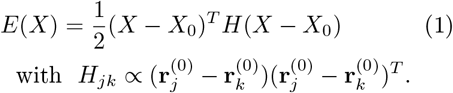

Solving the equations of motion in *E* leads to the eigenvalue problem *HU* = Λ*U* where each of the 3*M* eigenpairs (*λ*_*l*_, *U*_*l*_) defines a *normal mode* of frequency 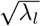 that deforms the reference conformation along 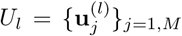. Pragmatically, the normal modes provide a linear basis of functions that spans the whole conformational space. Because the variance of each mode scales with the inverse of its frequency, it is customary to perform a low-rank approximation of *H* by only considering the *d* lowest frequency normal modes 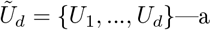 choice empirically justified as they often encode relevant functional motions [17, 37]. Formally, for rank *d*, the reference atomic model can be deformed as follows:

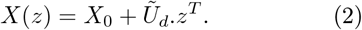

In practice, the eigenvalue problem is challenging to solve for large *M* as the Hessian is a 3*M* ×3*M* matrix. Because the lowest-frequency modes usually exhibit strong collectivity [17], *H*_*CG*_ is usually computed for a subset of atoms, the *coarse-grained (CG) model*. Its eigenvectors can then be interpolated on the remaining set of atoms and orthonormalized to yield 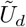.

#### Isolated Atom Superposition Approximation, Weak-phase Approximation and Projection Assumption

The electrostatic potential of the whole molecule can be approximated as the superposition of the electron form factors of its constitutive atoms [38]. Each form factor is defined as the sum of 5 Gaussians—their amplitude *a* and width *b* determined by the atomic type [39]. For typical cryo-EM experiments, the electron wave scattered by the atoms of the sample, which ultimately forms an image on a detector, can be linearized and simplifies to the integral of the volume, after it has been rotated by *R*_*i*_, along the path of the probing electron [40], resulting in the following “pre-image” *I*_*i*_, i.e. a mapping from ℝ^2^ toℝ:

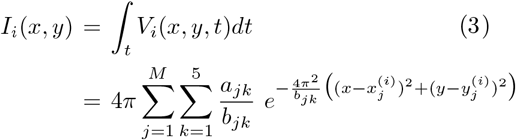

where 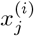 and 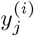 are the planar coordinates of atom *j* in volume *i* derived from 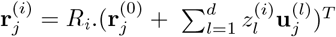.

The interaction between the beam and the microscope’s lens is modeled by the Point Spread Function (PSF) *P*_*i*_. Imperfect centering of the molecule in the image is characterized by small translations **t**_*i*_*∈*ℝ ^2^. The translation **t**_*i*_ and orientation *R*_*i*_ of the molecule are jointly referred to as its pose. Finally, taking into account signal arising from the vitreous ice in which the molecules are embedded as well as the non-idealities of the lens and the detector, each image *Ŷ*_*i*_ can be modelled as [38, 41]

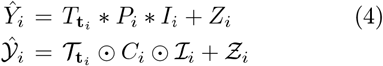

where * is the convolution operator, ⊙ indicates element-wise multiplication, *T*_**t**_ is the **t**-translation kernel and *Z*_*i*_ white Gaussian noise on ℝ^2^. In Fourier space, we note 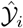 the Fourier transform of *Ŷ*_*i*_,ℐ _*i*_ the complex Fourier transform of the ideal image *I*_*i*_, 𝒯_**t**_ the **t**-translation operator or phaseshift in Fourier space, *C*_*i*_ is the Contrast Transfer Function (CTF), Fourier transform of the PSF, and 𝒵_*i*_ the Fourier transform of *Z*_*i*_.

## Datasets

### EMPIAR-10028

This cryo-EM dataset images the emitine bound *P*.*falciparum* 80S ribosome [27] (*Pf* 80S). This dataset consists of 105,247 images that were first processed with CryoSPARC [42] to find per-particle pose and CTF parameters. Particle images were downsampled to 128 × 128 pixels and fed to `aNiMAte` starting from an initial atomic model consisting of the two published structures of the *Pf* 80S ribosome small subunit (PDB 3J7A) and the large subunit (PDB 3J79).

#### EMPIAR-10180

This dataset [26] is known to contain both compositional and continuous heterogeneity. Compositional heterogeneity occurs when imaged particles are not structurally consistent across images of the dataset. It is worth noting that our method is not currently able to deal with compositional heterogeneity, mainly because of the use of a reference atomic model that is kept structurally constant during training. For this reason we selected 139,722 particle images that are believed to contain intact pre-catalytic spliceosome particles from the deposited EMPIAR-10180 dataset as described in the cryoDRGN paper [29]. Images were downsampled to 128 × 128 pixels and the atomic model associated with this dataset (PDB 5NRL) was used as a reference atomic model.

### Overview of the Architecture

#### Convolutional Encoder

The encoder learns a latent embedding of each image by mapping it to an estimate of the conformation variable *z*_*i*_ associated with it. The encoder is structured sequentially as a Convolutional Neural Network (CNN) containing 6 blocks, each consisting of 2 *Conv3* × *3* layers, followed by batch normalization and average pooling by a factor of 2.

The filter numbers for the convolutional layers in the blocks are set as [32, 64, 128, 256, 512, 1024]. The architecture of the CNN is inspired from the first layers of VGG16 [43], known to perform well on visual tasks. A *conformation* Multilayer Perceptron (MLP) with 2 hidden layers of sizes [512, 256] that maps to *z*_*i*_ of dimension *d*, which is the number of normal modes used.

#### Diffierentiable Simulation Decoder

The decoder is implemented as a simulation of the image formation model. Unlike the encoder, it is not parameterized by a neural network.

##### Pre-processing

A reference atomic model *X*_0_ is fed to ProDy [44] to pre-compute the Hessian matrix *H*_*CG*_ of the coarse-grained (CG) model. *H*_*CG*_ is diagonalized on the GPU [45] and its first *d* eigenvectors are interpolated on the remaining set of atoms with PyKeOps [46] to yield 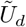.

The electron form factors [39] are retrieved with Gemmi [47]. For each image *i*, the rotation matrix, image shift and CTF are provided either by the simulator or by an external software like RELION [41] or cryoSPARC [48].

##### Training

The forward pass is differentiable with respect to *z*_*i*_. For each image 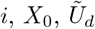 and *z*_*i*_ are combined according to Eq.2. The resulting atomic coordinates are rotated by *R*_*i*_ *via* a matrix multiplication. A fixed grid of *D*^2^ coordinates on the x-y plane in real space is used to evaluate Eq.3. The resulting image is Fourier transformed and element-wise multiplied by 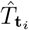 and *C*_*i*_. A final inverse Fourier transform step is carried out.

#### Training Procedure

The training procedure for both experimental datasets presented here follows the pipeline described above. For every training run, an initial automatic alignment step is performed to ensure the reference atomic model is in the same coordinate system as the acquired particle images (consensus reconstruction). This alignment step relies on the same architecture presented in Fig. 1 (A), where the encoder *ψ* is frozen and a global pose, parameterized as a 4 x00D7; 4 transformation matrix, for *X*_0_ is searched through stochastic gradient descent and automatic differentiation of the decoder. After the initial alignment step is done, the encoder is unfrozen and the whole architecture with the global pose is jointly optimized. A normalized crosscorrelation loss ℒ is the objective used in training. This statistical loss removes the need for finding a normalization constant to account for intensity discrepancy between the reconstructed 2D image from Eq.3 and the experimental cryo-EM image.

Training was carried with the following parameters: we use the Adam optimizer [49] with a learning rate of 1 × 10^−3^ for the alignment stage, and 1 × 10^−4^ for the joint optimization, a minibatch size of 32. Alignment was run for the first 10,000 minibatch steps, with the remaining time dedicated the joint optimization of encoder and global pose. Each training run was stopped after 24 hours (approximately 65 epochs for EMPIAR-10028 and 50 epochs for EMPIAR-10180). The whole framework is implemented in PyTorch and is run on 16 nVidia Geforce 2080Ti GPUs through distributed data parallel training.

## Contributions

Y.N, J.M. and F.P. conceived the idea of `aNiMAte`. Y.N. and D.R. conceived the scoring feature. F.P. and D.R. supervised the research. Y.N. and J.M. implemented the code with contributions from A.P., A.L., H.M., and F.P. Y.N. conducted all the experiments. G.W., N.M. provided critical guidance to Y.N., J.M. and F.P. F.P. wrote the paper with input from all the authors.

## Data Availability

The spliceosome and ribosome datasets are publicly available from the Electron Microscopy Public Image Archive with IDs 10180 [26] and 10028 [27].

## Code Availability

The source code of `aNiMAte` is available on GitHub at github.com/compSPI/aNiMAte.

## Notes

### Competing Interest Statement

The authors have declared no competing interest.

### Funding

- United States Department of Energy: DE-AC02-76SF00515, the LCLS seed grant "From Atomic Models to Noisy Images and Back Again: End-to-end Differentiable SPI simulators for CryoEM and XFELs" (PI: G.W.)
- SLAC National Accelerator Laboratory: LDRD project "AtomicSPI: Learning atomic scale biomolecular dynamics from single-particle imaging data" (PI: F.P., Y.N.)
- SLAC National Accelerator Laboratory: Machine Learning Initiative
- United States Department of Health and Human Services: National Institutes of Health (NIH) grant No. 1R01GM144965-02

