## Supplementary Figures for "Unsupervised learning of structural variability in cryo-EM data using normal mode analysis of deformable atomic models"

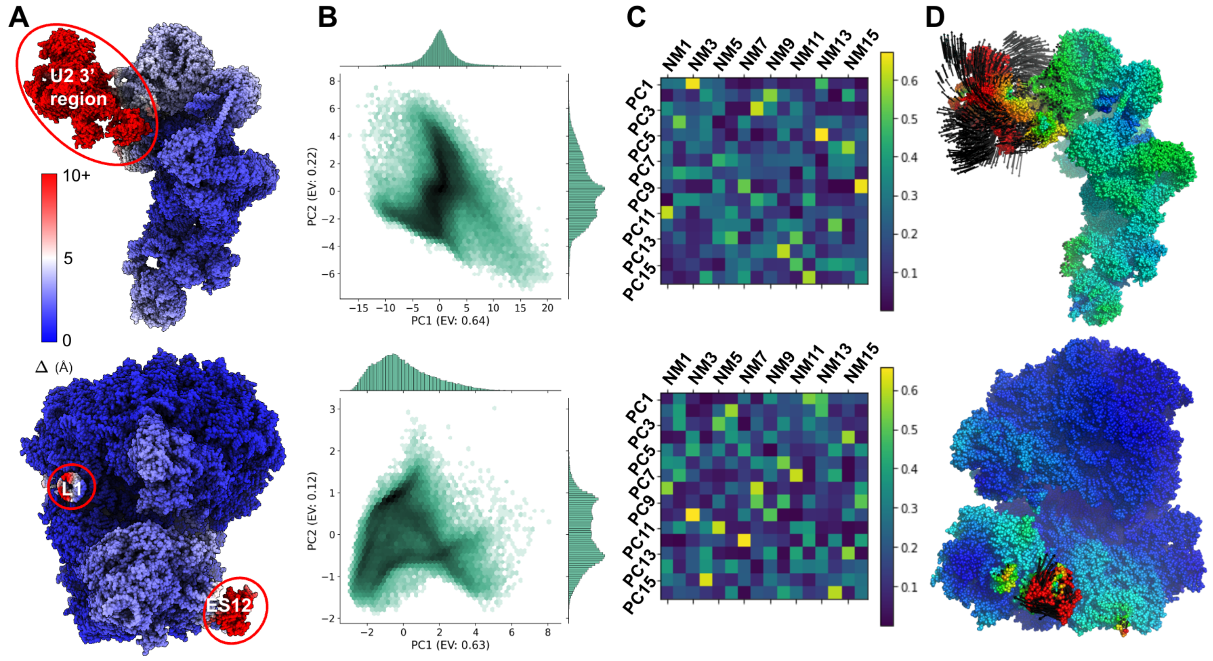


**Supplemental Material, Figure S1 Results on experiment datasets with deposited atomic models: top - EMPIAR 10180 (spliceosome); bottom - EMPIAR 10028 (ribosome).** **(A)** The atomic model used to train `aNiMAte` is shown colored with its atomic model score Δ. Regions with atomic model scores above 10Å are highlighted. These regions are discarded to produce the Δ-reduced atomic models used to train `aNiMAte` as presented in the main text. **(B)** Principal Component Analysis of the learned normal mode coordinates results in a 2D space (PC1, PC2) explaining more than 75% of the dataset variability. **(C)** The relative contribution of the 16 normal modes computed from the atomic model is shown for each principal component. **(D)** The first principal component for each dataset is illustrated.

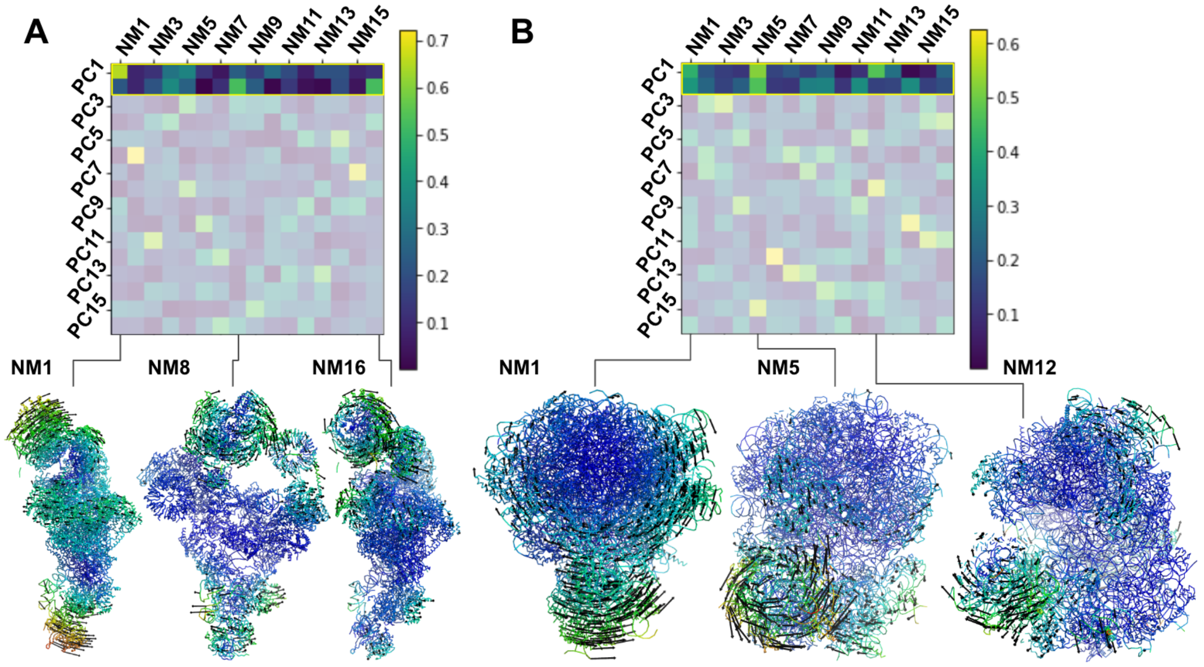


**Supplemental Material, Figure S2 Results on experiment datasets with Δ-reduced atomic models: A. EMPIAR 10180 (spliceosome), B. EMPIAR 10028 (ribosome)**. For each dataset, the heatmap represents the contribution of each pre-computed normal mode (NM) to the predicted principal components (PC). The first two PCs are highlighted, and the main three NMs contributing to them are further illustrated in the space of atomic models.
